# Residual force enhancement is not altered while force depression is amplified at the cellular level in old age

**DOI:** 10.1101/2024.06.14.599072

**Authors:** Binta S. Njai, Avery Hinks, Makenna A. Patterson, Geoffrey A. Power

**Author notes:** Correspondence:* Geoffrey A Power PhD. **N**euromechanical **P**erformance **R**esearch **L**aboratory, Department of Human Health and Nutritional Sciences, College of Biological Sciences, University of Guelph, Ontario, Canada.

## Abstract

Residual force enhancement (rFE) and residual force depression (rFD) are history-dependent properties of muscle which refer to increased and decreased isometric force following a lengthening or shortening contraction, respectively. The history-dependence of force is greater in older than younger adults when assessed at the joint level. However, it is unclear whether this amplification of the history-dependence of force in old age is owing to cellular mechanisms or a consequence of age-related remodeling of muscle architecture. Single muscle fibres from the psoas major of old and young F344BN rats were dissected and chemically permeabilized. Single muscle fibres were mounted between a force transducer and length controller, then maximally activated (pCa 4.5). To assess rFD, fibers were actively shortened from 3.1 to 2.5µm at both a slow (0.15Lo/s) and fast (0.6Lo/s) speed, with a fixed-end isometric reference contraction at 2.5µm. To assess rFE, fibers were activated and stretched at 0.3Lo/s from a sarcomere length of 2.2 to 2.5µm, and 2.7 to 3.0µm, and compared to fixed-end isometric reference contractions at 2.5 and 3.0µm, respectively. Isometric force was ≈19% lower in old as compared with young (p<0.001). Upon normalizing to fibre cross-sectional area, there was no age-related difference in specific force (p>0.05). rFD was ≈33% greater in old as compared with young (p<0.05), while rFE did not differ between groups (p>0.05). rFD is amplified in old age at the cellular level, while rFE appears to be unchanged, thus previously reported age-related modification of rFE occurs upstream from the cellular level.

## Introduction

The history-dependent properties of muscle known as residual force enhancement (rFE) and residual force depression (rFD) refer to an increase and decrease in isometric force following a lengthening (i.e., eccentric) or shortening (i.e., concentric) contraction, respectively, as compared to a fixed-end isometric contraction at the same muscle length and level of activation (Abbott and Aubert, 1952; Hahn et al., 2023; Seiberl et al., 2015). The presence of rFE and rFD has been observed across all scales of muscle (Chen et al., 2020; Joumaa et al., 2008; Leonard et al., 2010; Mashouri et al., 2021; Pinnell et al., 2019; Ramsey et al., 2010), and at the joint level in humans during electrically stimulated and voluntary contractions (Chapman et al., 2018; Chen et al., 2019; Hahn et al., 2023; Seiberl et al., 2015). Both rFE and rFD appear to be greater in older as compared with younger adults when assessed at the joint level during voluntary contractions (Power et al., 2012; Power et al., 2014a; Power et al., 2015). However, it is unclear whether this amplification of the history-dependence of force in old age is owing to cellular mechanisms or a consequence of age-related muscle architecture remodeling.

The mechanisms of rFE and rFD originate at the cellular level. For rFE, in the presence of Ca^2+^ and cross-bridge cycling, ‘the molecular spring’ titin becomes stiffer thereby contributing greater tension during and following active lengthening of the muscle (Hahn et al., 2023; Herzog, 2018; Hessel et al., 2023), resulting in a greater contribution of passive force to total force production as compared with a fix-end isometric contraction (Herzog and Leonard, 2002). As well, rFE is greatest following large stretch amplitudes to long muscle lengths (Bakenecker et al., 2022; Fukutani et al., 2017; Rassier and Herzog, 2004). rFD occurs due to an inhibition of cross-bridge attachment following active shortening of the muscle, limiting available binding sites, resulting in fewer attached force generating cross-bridges as compared to a fix-end isometric contraction (Hahn et al., 2023; Joumaa et al., 2017; Joumaa et al., 2021), and is proportional to work of shortening (Herzog et al., 2000).

Interestingly, both rFE and rFD were amplified in older compared to young adults at the joint level (Power et al., 2012; Power et al., 2014a; Power et al., 2015). This age-related amplification of the history-dependence of force may be explained by changes in muscle architecture, as muscle fascicle length decreases with age (Hinks et al., 2024; Narici et al., 2003; Power et al., 2013a; Power et al., 2021). Therefore, older individuals with shorter muscle fascicles may have experienced relatively greater changes in sarcomere length during joint-level active lengthening and shortening contractions, leading to greater rFE and rFD, respectively. Whether the amplification of the history-dependence of force in old age originates at the cellular level has not been investigated, leaving the mechanisms behind an amplification of rFE and rFD in old age unclear. Therefore, in the present study we investigated whether age-related differences in rFE and rFD exist at the single muscle fibre level.

## Methods

### Animals

Ten young (sacrificial age ∼8 months; mass = 381.61 ± 15.71 g) and fourteen old (sacrificial age ∼32 months; mass = 480.79 ± 37.91 g) male F344BN F1 rats were obtained from the National Institute on Aging aged rodent colonies (Charles River Laboratories, Senneville, QC, Canada) with approval from the Animal Care Committee (AUP: #4905) at the University of Guelph and all protocols following guidelines from the Canadian Council on Animal Care. Rats were housed in groups of two or three with a 12 h light/dark cycle at 23°C and were given unrestricted access to room-temperature water and a Teklad global 18% protein rodent diet (Envigo, Huntington, Cambridgeshire, UK). Rats were sacrificed via isoflurane followed by CO_2_ asphyxiation then cervical dislocation prior to harvesting muscle tissue.

### Tissue Preparation

The right and left psoas major muscles were harvested from the rats following sacrifice and immediately transferred to a silicone elastomer-plated petri dish containing chilled dissecting solution. The psoas major muscle was selected owing to its long fibres and almost complete fast-type fibre composition (Hämäläinen and Pette, 1993; Schilling et al., 2005; Vlahovic et al., 2017). Fibre bundles were dissected from the proximal half of the psoas major, which is almost exclusively fast-type fibres (< 1% type I) (Hämäläinen and Pette, 1993). The muscles were then sectioned into fibre bundles and transferred to a tube containing 2.5 ml of chilled skinning solution where they remained on ice for 30 min for permeabilization (Hubbard et al., 2023; Mazara et al., 2021). The bundles were then washed with fresh chilled dissecting solution and gently agitated to ensure any remaining skinning solution was removed. The bundles were then transferred to a tube containing storage solution and incubated for 24 hr at 4°C. Tubes were prepared with fresh storage solution and the bundles were placed in individual tubes and stored in a freezer at -80°C until mechanical testing began (Hubbard et al., 2023; Mashouri et al., 2021).

### Solutions

The dissecting solution was composed of the following (in mM): K-propionate (250), Imidazole (40), EGTA (10), MgCl_2_·6H_2_O (4), Na_2_H_2_ATP (2). The storage solution was composed of K-propionate (250), Imidazole (40), EGTA (10), MgCl_2_·6H_2_O (4), Na_2_H_2_ATP (2), glycerol (50% of total volume after transfer to 50:50 dissecting: glycerol solution). The skinning solution with Brij 58 was composed of K-propionate (250), Imidazole (40), EGTA (10), MgCl_2_·6H_2_O (4), 1 g of Brij 58 (0.5% w/v). The relaxing solution was composed of Imidazole (59.4), KMSA (86), Ca(MSA)_2_ (0.13), Mg(MSA)_2_ (10.8), K_3_EGTA (5.5), KH_2_PO_4_ (1), H_2_O, Leupeptin (0.05), Na_2_ATP (5.1). The pre-activating solution was composed of K-propionate (185), MOPS (20), Mg(CH_3_COO)_2_ (2.5), Na_2_ATP (2.5). The activating solution (pCa 4.5) was composed of Ca^2+^ (15.11), Mg (6.93), EGTA (15), MOPS (80), Na_2_ATP (5), CP (15). All solutions were adjusted to a pH of 7.0 with the appropriate acid (HCl) or base (KOH).

### Mechanical Testing and Force Measurements

On testing days, single muscle fibres were isolated from a fibre bundle and moved to a temperature-controlled chamber filled with relaxing solution where they were then tied to pins connected to a force transducer (403A, Aurora Scientific, Toronto, ON, Canada) and a length controller (322C, Aurora Scientific) with nylon sutures. All experiments were performed at 12°C to allow for a balance between force production capacity, and the number of contractions performed before experiencing force loss (Bottinelli et al., 1996; Ranatunga, 1982; Ranatunga, 1984; Tomalka et al., 2017; Tomalka et al., 2021). Muscle contraction was initiated by first transferring the fibre from relaxing solution to a pre-activating solution with reduced Ca^2+^ buffering capacity for 10 s before being transferred to an activating solution. All activations were performed in a maximal Ca^2+^ concentration solution (pCa 4.5). For all tests, the average sarcomere length (SL) was measured using a high-speed camera (Aurora Scientific Inc., HVSL 901A). Total force was recorded, and active force was determined by subtracting passive from total force. Before starting the testing protocol, the fibre was set to a SL of 2.5 μm (i.e., the optimal SL for rat muscle fibres (Burkholder and Lieber, 2001; Ledvina and Segal, 1995)) and a ‘fitness’ contraction was performed to ensure the ties were not loose and the condition of the fibre was sufficient for testing. After the fitness test, SL was re-measured and, if necessary, re-adjusted to 2.5 μm. Then, the cross-sectional area (CSA) of each fibre was determined. This was done by measuring the diameter of the fibre in three places (at each end and in the centre) using an eyepiece reticle. These three measurements were then averaged to calculate the CSA of the individual fibre. Given these samples are muscle fibre segments and chemically permeabilized, they are treated as cylindrical, therefore we assumed circularity in the calculation of CSA.

### Force Depression

Fibres were set to an average SL of 2.5 μm, then passively lengthened to 3.1μm. Activation began 10 s after lengthening to 3.1 µm. After 20 s of activation, fibres were actively shortened to an average SL of 2.5 μm at 0.15 L_O_/s (slow speed; Figure 1A) and held for 15 s to reach an isometric steady-state before deactivation. The same protocol was repeated with a velocity of 0.6 L_O_/s (fast speed; Figure 1B), with the order randomized between the two velocities. For the isometric reference contraction (ISO), fibres were activated at an average SL of 2.5 μm for 35 s.

**Figure 1:**
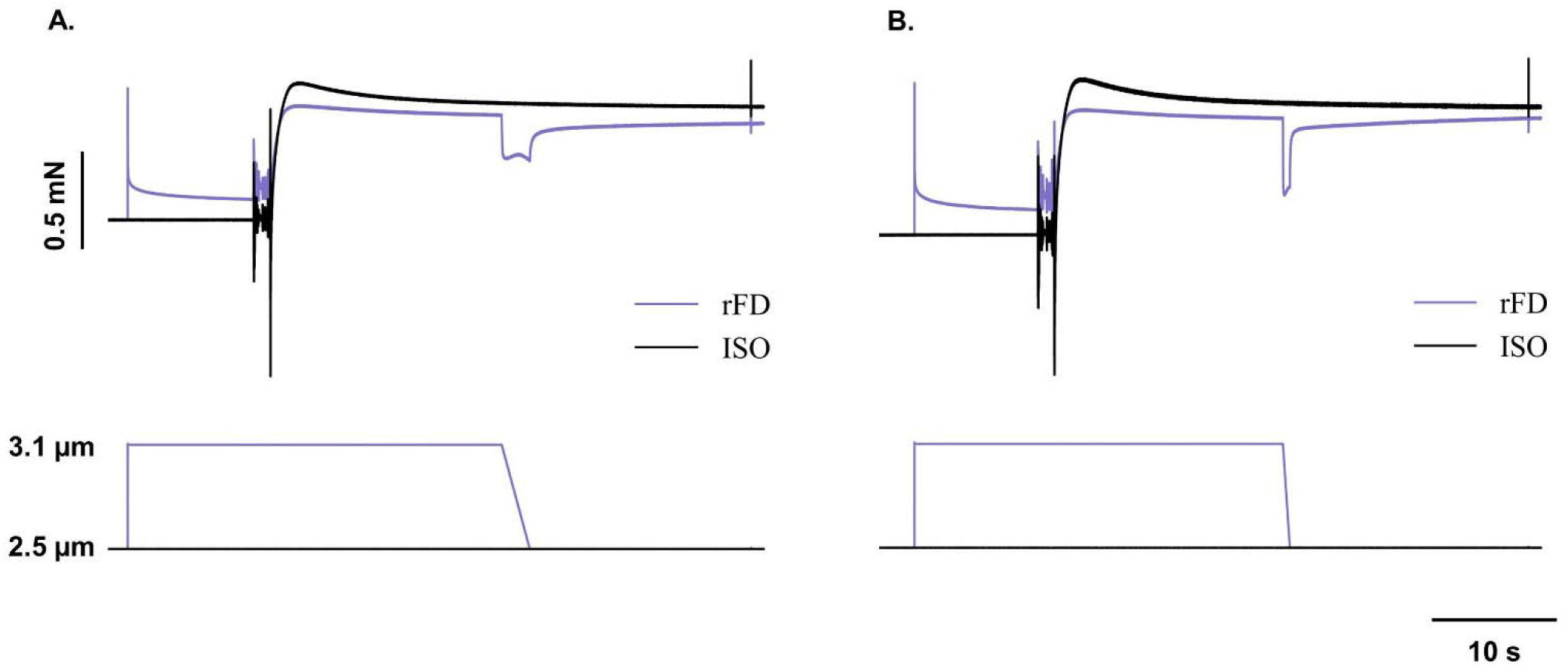
Representative force trace for residual force depression (rFD; purple) and isometric reference (ISO; black) conditions. (**A.**) rFD passively lengthened to 3.1 µm, activated and shortened to 2.5 µm (i.e., plateau of the force-length relationship) at 0.15 L_O_/s, representative of a slow velocity shortening contraction. (**B.**) rFD protocol repeated at the same lengths with the exception of increased velocity of 0.6 L_O_/s. An instantaneous stiffness test was performed in both conditions prior to deactivation.

### Residual force enhancement

For assessment of rFE near the plateau-region of the force-length relationship, first, a reference isometric contraction was performed at a SL of 2.5 µm. For the stretch condition, fibres were set to an average SL of 2.5 µm and passively shortened to 2.2 µm, then 10 s following passive shortening were activated. After 20 s of activation, fibres were actively stretched to 2.5 µm at 0.3 L_O_/s and held for 15 s before deactivation (Figure 2A). For assessment of rFE on the descending limb of the force-length relationship, a reference isometric contraction was performed at a SL of 3.0 µm. Fibres were set to an average SL of 3.0 µm and passively shortened to 2.7 µm. Activation began 10 s after passive shortening. After 20 s of activation, fibres were actively stretched to 3.0 µm at 0.3 L_O_/s and held for 15 s to reach an isometric steady-state before deactivation (Figure 2B).

**Figure 2:**
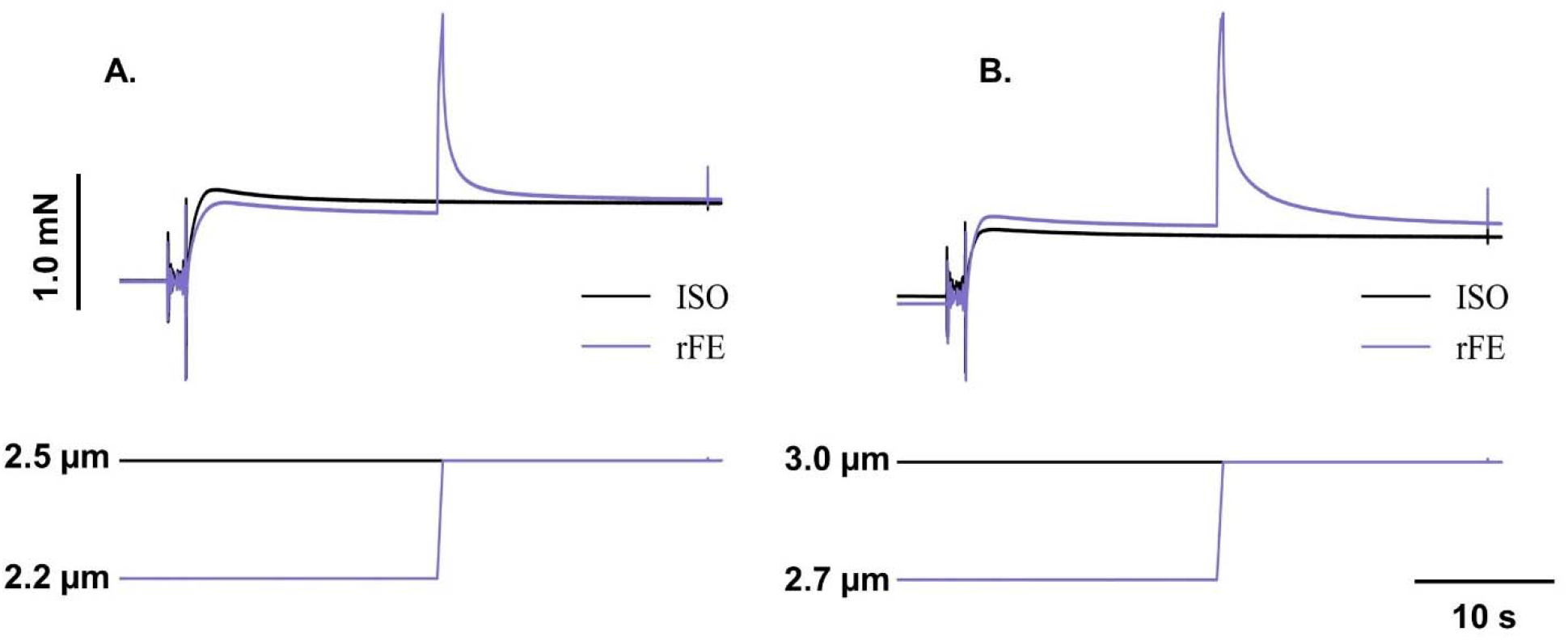
Representative force trace from a young single muscle fibre for residual force enhancement (rFE; purple) and isometric reference (ISO; black) conditions. (**A.**) rFE fibres set to average SL 2.2µm and passively stretched to 2.5µm (i.e., plateau of the force-length relationship) at 0.3 L_O_/s. Representative force tracing for force enhancement (purple) and isometric reference (black) conditions. (**B.**) Fibres set to average SL of 2.7 µm and passively stretched to 3.0 µm at 0.3 L_O_/s (i.e., descending limb of the force-length relationship).

### Analysis

Based on previous protocols from our lab (Patterson et al., 2024), to ensure fibre fidelity, we excluded any fibres that produced isometric active specific force less than 20 mN/mm^2^. Furthermore, we removed any fibres from analysis that experienced greater than a 20% drop in force from the first ISO contraction of the rFD protocols to the ISO contraction of the short rFE protocol (both were at a sarcomere length of 2.5 µm). Next, the force traces of any fibres that exhibited positive rFD (i.e., greater steady state force in the rFD contraction compared to the ISO contraction) were manually inspected for any signs that the fibre ripped or other errors in the signal. If these issues were observed, those fibres were removed from analysis. We manually inspected the force traces of any fibres that exhibited negative rFE (i.e., lower steady state force in the rFE contraction compared to the ISO contraction) for any signs of ripping or eccentric contraction-induced damage. If damage-induced force loss was evident, those fibres were removed from analysis. Lastly, because more rFE than rFD protocols were removed from the initial analyses, additional fibres were tested for only the rFE protocols to ensure sample sizes among all protocols were balanced. Altogether, these data inclusion steps left us with a final sample size of 40 old and 49 young fibres for the fast rFD protocol, 42 old and 48 young fibres for the slow rFD protocol, 49 old and 54 young fibres for the short rFE protocol, and 43 old and 48 young fibres for the long rFE protocol.

### Work of shortening

Mechanical work of shortening during the rFD protocols was calculated as the product of the area under the curve of the force-time trace during shortening and the change in fibre length.

### Stiffness

Instantaneous stiffness (k) tests were performed to determine the proportion of attached cross-bridges during the force plateau. This was done by rapidly stretching the fibre (500 L_O_/s) by 0.3% of L_O_/s and dividing the change in force during the stretch by the change in length.

### History-dependent properties of force

For rFD, rFE, and the ISO values, force was reported as an average over 500 ms before the instantaneous stiffness test at the same time following activation for all conditions (Figures 1 & 2). For the rFE and rFD trials, the 500 ms time point corresponded to the isometric steady-state 15 s following the dissipation of lengthening/shortening force transients. rFD was calculated as the difference between the average force in the rFD condition compared to the ISO condition at 2.5 μm. Similarly, stiffness depression was calculated as the % decrease in instantaneous stiffness in the rFD state compared to ISO (Joumaa et al., 2012). rFE was calculated as the difference between the force in the rFE condition compared to the ISO condition at 2.5 (short) and 3.0 μm (long).

All statistical analyses were performed in SPSS Statistics Premium 28. Age-related differences in fibre CSA, maximum isometric force at 2.5 μm, and instantaneous stiffness were assessed by one-way analysis of variance (ANOVA). Age- and speed-related differences in rFD, stiffness depression, and work of shortening were assessed via two-way ANOVA (speed [fast, slow] × age [young, old]). Age- and length-related differences in rFE were assessed by a two-way ANOVA (length [short, long] × age [young, old]). To assess the relationship between %rFD and %stiffness depression, a linear regression was performed between %rFD and the change in instantaneous stiffness between the ISO and the history-dependent conditions. Data are reported in figures as the mean ± standard deviation, α=0.05.

## Results

Single muscle fibres from old were ≈19% weaker (p=0.002; Figure 3A.), with an ≈19% smaller CSA (p=0.001; Figure 3B.) and ≈48% lower (p<0.001) instantaneous stiffness at maximal activation (i.e., a smaller proportion of attached cross-bridges in old; Figure 3C.) as compared with young. When force was normalized to CSA to account for the lower myofibrillar protein content in old, there was no age-related difference (p=0.807) in specific force (Figure 3D.), indicating intrinsic muscle quality was intact.

**Figure 3:**
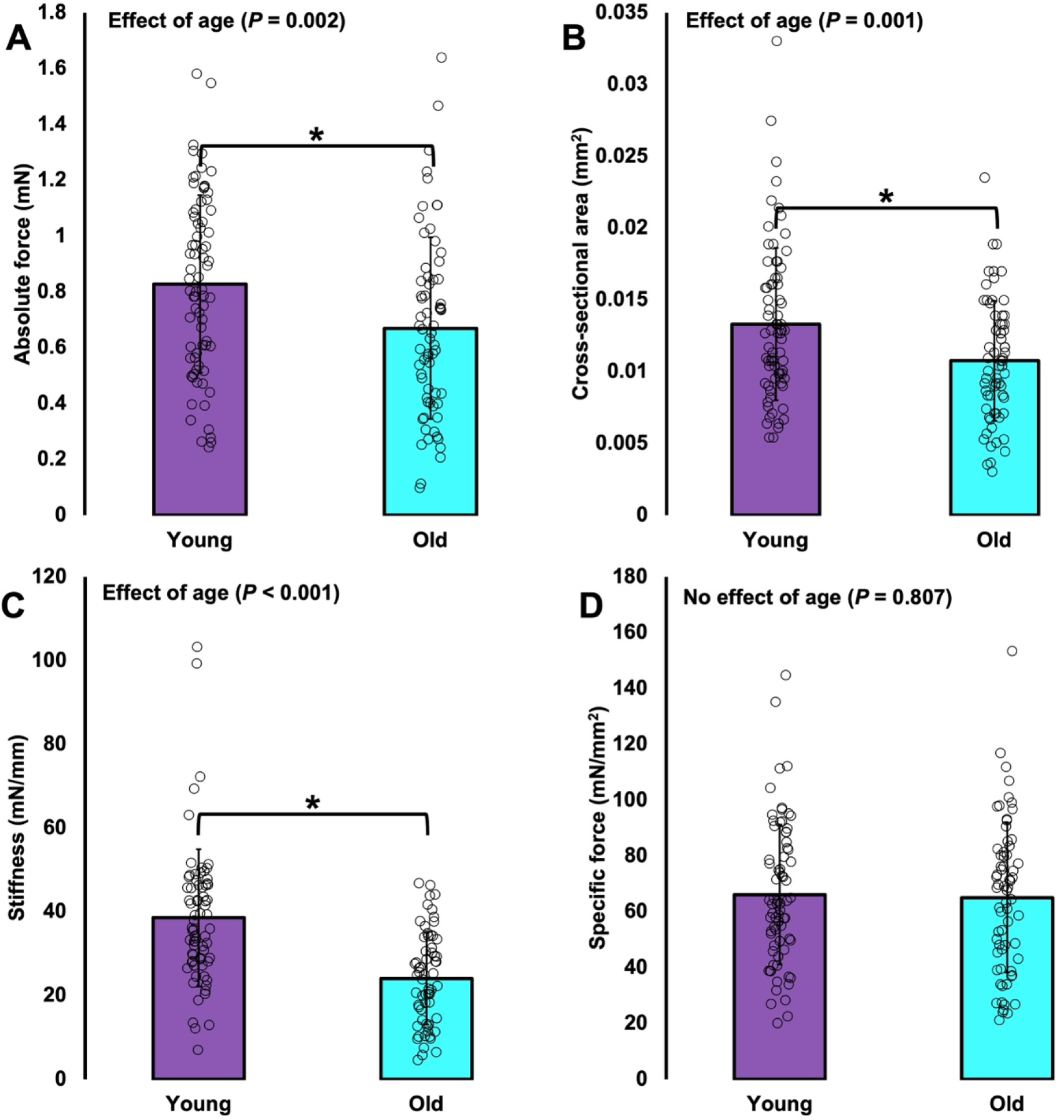
Single muscle fibre cross sectional area and isometric force properties. Absolute force (**A.**), cross-sectional area (**B.**; CSA), and instantaneous stiffness (**C.**) was lower in old as compared with young. When force was normalized to CSA, there was no age-related difference in specific force (**D.**). **\***Difference between young and old (p<0.05).

### Residual force enhancement

For absolute rFE (mN) there was no interaction of age × length (p=0.454), nor effect of age (p=0.575), but as expected, rFE was ≈109% greater on the descending limb (sarcomere length: 3.0 µm) as compared with the plateau region (sarcomere length: 3.0 µm) of the force-length relationship (p<0.001; Figure 4A.). For relative rFE (%) there was no interaction of age × length (p=1.00), nor effect of age (p=0.229), and like absolute rFE, relative rFE was ≈135% greater on the descending limb (long) as compared with the plateau region (short) of the force-length relationship (p<0.001; Figure 4B.).

**Figure 4:**
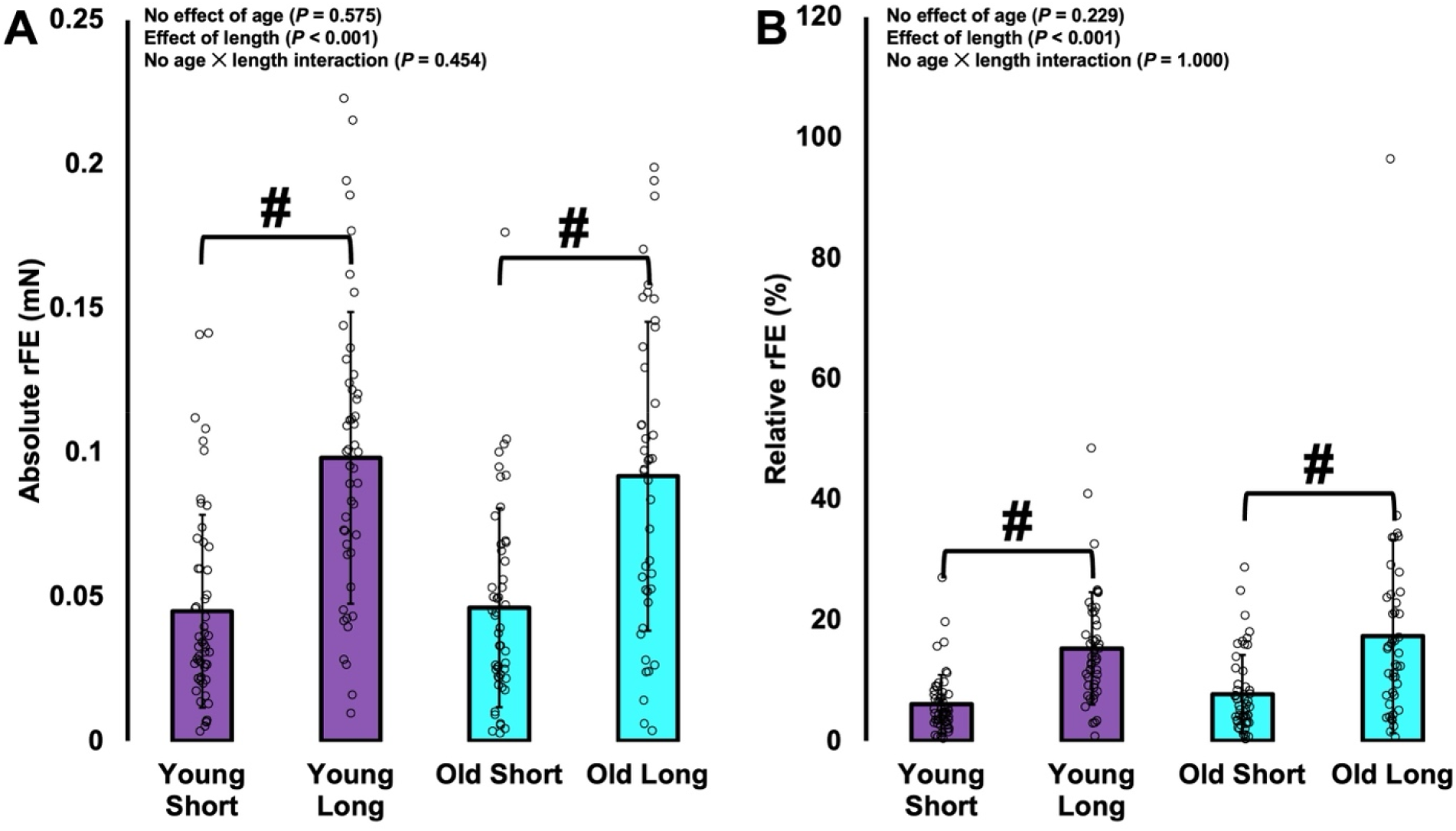
Absolute (**A.**) and relative force enhancement (rFE) (**B.**), on the plateau (short; sarcomere length: 2.5µm) and descending limb (long; sarcomere length: 3.0 µm) of the force-length relationship. rFE was greater at longer sarcomere lengths as compared with shorter, with no age-related differences. **#**Difference between short and long with young and old combined (p<0.05).

### Work of shortening

For work performed during the active shortening contractions, there was no interaction of age × speed (p=0.228), but there were main effects for both age (p=0.007) and speed of shortening (p<0.001), such that old performed ≈26% less work than young, and ≈110% more work was performed during the slow as compared with the fast speed condition (Figure 5D.).

**Figure 5:**
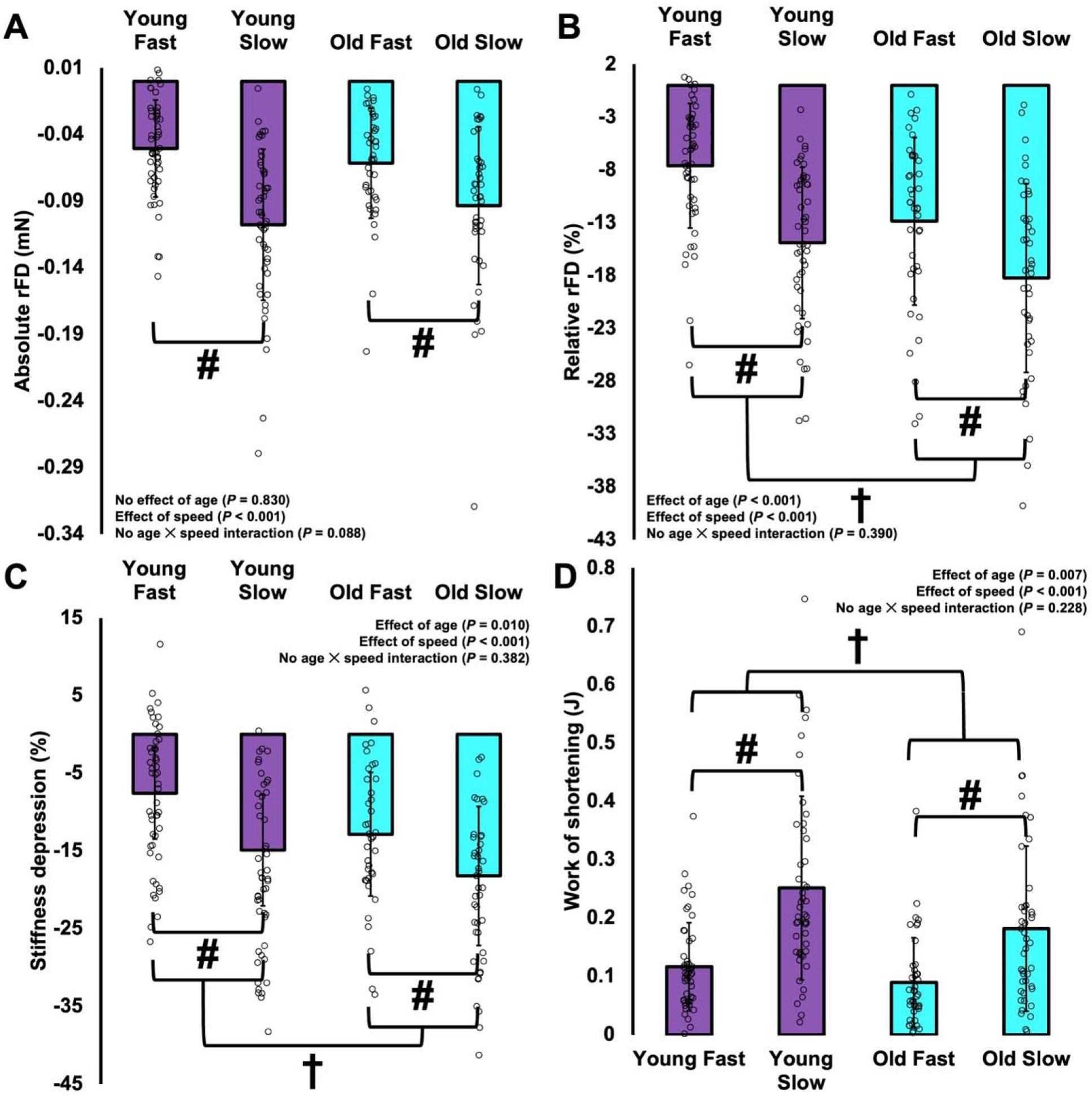
Absolute (**A.**) and relative force depression (rFD) (**B.**), % stiffness depression (**C.**), and work (**D.**) in young and old when shortening from an average sarcomere length of 3.1 to 2.5 µm at a fast (0.60Lo/s) and slow speed (0.15Lo/s). Both absolute and relative rFD was greater in the slow than fast condition, and old experienced greater rFD(%) as compared with young. There was a greater reduction in stiffness in the slow speed condition, and old had greater stiffness depression as compared with young. Work was greater during the slow as compared with fast condition, and lower in old as compared with young. **#**Difference between slow and fast with young and old combined (p<0.05). **†**Difference between young and old with slow and fast combined (p<0.05).

### Residual force depression

For absolute rFD (mN) there was no interaction of age × speed (p=0.088), nor effect of age (p=0.830), but as expected, rFD was ≈80% greater for the slow as compared with fast speed condition (p<0.001; Figure 5A.), owing to greater work of shortening during the slow as compared with the fast condition. For relative rFD (%) there was no interaction of age × speed (p=0.390), however there were main effects for both age (p<0.001) and speed (p<0.001), such that old had ≈38% greater rFD (%) as compared with young despite performing less mechanical work of shortening, and there was ≈62% more rFD (%) for the slow as compared with fast condition (Figure 5B.).

### Stiffness

For instantaneous stiffness depression (%) there was no interaction of age × speed (p=0.382), but there was an effect of speed with ≈62% greater reductions in stiffness for the slow as compared with fast condition (p<0.001). As well, there was ≈38% greater stiffness depression in old compared to young (p=0.010; Figure 5C.), notably this was the same magnitude of age-related difference as for relative rFD.

### Linear regression analysis

For both the fast (Figure 6A.) and slow (Figure 6B.) shortening speed conditions, in both young and old there were strong relationships (R^2^ = 0.59-0.84, all p<0.001) between rFD (%) and stiffness depression (%). These relationships tended to be slightly weaker in old (R^2^=0.59-0.65) as compared with young (R^2^=0.78-0.84), such that stiffness depression did not explain as much of the variance in rFD in old as in young, indicating that the greater rFD in old may be driven by additional factors outside of fewer attached cross-bridges.

**Figure 6:**
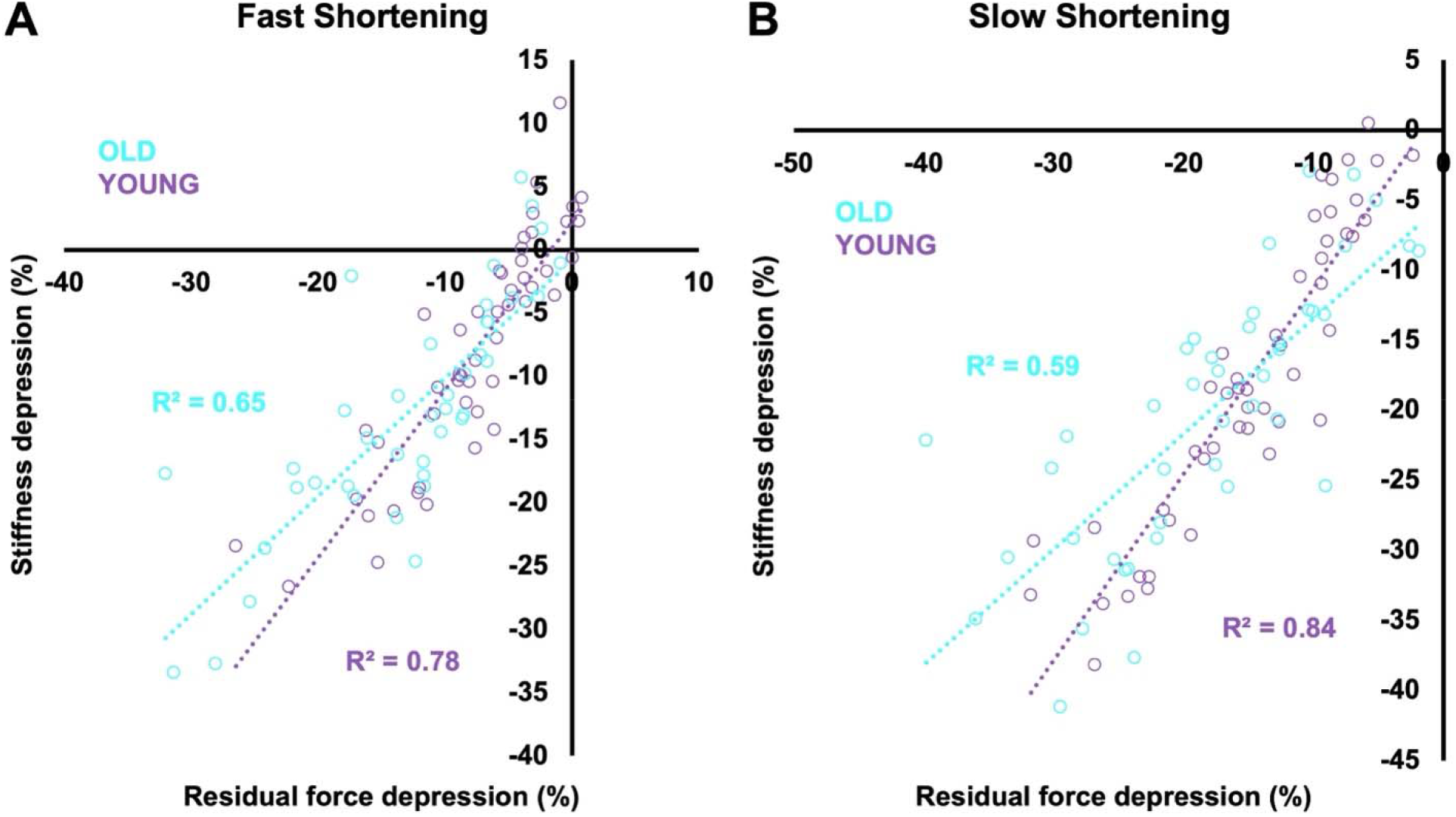
Linear regression analysis of stiffness depression(%) versus rFD(%) for the slow and fast condition in young (black circles) and old (white circles). (**A.**) Fast condition: combined groups (solid line, R^2^=0.73, p<0.05), Young group (dotted line, R^2^=0.78, p<0.05), Old group (dashed line, R^2^=0.68, p<0.05). (**B.**) Slow condition: combined groups (solid line, R^2^=0.70, p<0.05), Young group (dotted line, R^2^=0.84, p<0.05), Old group (dashed line, R^2^=0.63, p<0.05),

## Discussion

The history-dependence of force is an intrinsic property of muscle currently unexplained by the cross-bridge theory. In the present study we show that the amplification of rFD in old age originates at the cellular level and is indeed an intrinsic property of muscle which scales to joint-level voluntary contractions. Meanwhile, the cellular mechanisms contributing to rFE in old age are not altered, therefore previous reports of greater rFE in old age at the joint level are driven by factors upstream of the single muscle fibre.

### Age-related impairments in force production

*S*ingle muscle fibres from old rats exhibited typical age-related declines in maximal isometric force production, cross-sectional area, and the proportion of attached cross-bridges (Lim and Frontera, 2022; Lim and Frontera, 2023; Mazara et al., 2021). However, intrinsic contractility of the muscle appears to be intact as specific force (i.e., muscle quality) was not different between old and young when force was normalized to CSA accounting for the age-related decrease in myofibrillar protein content (Figure 3).

### History-dependence of force

In the present study our values of rFE at short and long lengths averaged 7% and 16%, respectively. As well, rFD values for the slow and fast shortening condition averaged -16% and -10%, respectively. These values are consistent with data published previously for single muscle fibres from the human vastus lateralis (0-19%; rFD, 6-25%; rFE (Pinnell et al., 2019)), and rabbit psoas and soleus (1-29%; rFD, (Joumaa et al., 2015)). Some of our rFD values were low or greater than 0%; our conservative approach in testing order may have contributed, whereby performing rFD first before the reference fixed-end isometric contraction could have lowered the force value recorded. Although, consistent with any of the low rFD values is a similar lack of stiffness depression (Figure 6), indicating that inhibition of cross-bridge attachment following active shortening did not occur in those fibres. While fibre type was not assessed, these values may represent some of the slowest fibres, unable to perform sufficient work of shortening during the fast condition inducing negligible rFD. While rFE appears to be velocity-independent (Herzog, 2004), rFD on the other hand is dependent on the work of shortening, and therefore is greater during slow contractile speeds where the muscle can generate higher forces than fast speeds (Herzog, 2004). We observed this velocity-dependence for rFD in the present study in single fibres from both young and old (Figure 5). Muscle fibre type has been suggested to influence the history-dependence of force with fast-type muscle having greater magnitudes of rFE (Ramsey et al., 2010) and rFD (Joumaa et al., 2015). However, others have not observed a fibre-type dependence for rFE (Pinnell et al., 2019). Given rFD is dependent on work of shortening, fast-type muscles experience greater rFD for a given speed of shortening as compared with slow-type. Upon normalizing to maximal shortening velocity, rFD does not appear to differ across fibre types (Joumaa et al., 2015). We tested muscle fibres from the proximal portion of the psoas major which is composed almost exclusively of type II fibres (Hämäläinen and Pette, 1993; Schilling et al., 2005; Vlahovic et al., 2017). It is unlikely that any differences in fibre type between young and old influenced our findings, as aging is typically associated with a greater proportion of slow-type fibres (Hepple and Rice, 2016; Kanda and Hashizume, 1989), and if old had more slow type fibres in the present study, they would have likely experienced less (rather than more) rFD than young.

### rFE in old age

As expected, there was no difference in the magnitude of rFE between young and old. Furthermore, rFE was indeed greater on descending limb than the plateau region of the force-length relationship (Edman et al., 1978; Rassier and Herzog, 2004) (Figure 4).

rFE in older adults has previously been reported to be ≈2.5 times higher than in young for the ankle dorsiflexors (Power et al., 2012), while the knee extensors had a similar level of RFE compared with young, but the relative contribution of passive force to total force production was greater in the older adults (Power et al., 2013a). As well, the time to reach an isometric steady-state force level following active lengthening was reported to be longer in old when compared with young (Power et al., 2014b). Therefore, we have previously hypothesised that a passive structural mechanism may play a disproportionately greater role in rFE in old compared to young (Power et al., 2012). However, given the absence of any age-related amplification of rFE at the cellular level in the present study (Figure 4), we disprove our original working hypothesis. Therefore, greater rFE magnitudes in older adults at the joint level are driven by factors upstream of the cellular mechanisms of rFE. A plausible explanation is a greater relative excursion of muscle fascicles for a given joint angular rotation, owing to muscle architectural remodeling resulting in shorter fascicle lengths in old age (Hinks et al., 2024; Narici et al., 2003; Power et al., 2013a; Power et al., 2021). With shorter fascicles than young, muscles of older adults would be lengthened relatively more, and may fall further along the descending limb of the force– length-relationship than those of young adults, and would be consistent with the finding that rFE is greatest following large stretch amplitudes to long muscle lengths (Bakenecker et al., 2022; Fukutani et al., 2017; Rassier and Herzog, 2004).

### Age-related amplification of rFD

rFD was greater in old as compared with young rat single fibres, and this finding aligns with our previous reports at the joint level in humans where rFD was greater during ankle dorsiflexion for older as compared to younger men (Power et al., 2014a). Additionally, in the present study the magnitude of rFD was greater during the slow shortening speed condition as compared with fast, likely owing to the greater work of shortening performed in the slow condition (Figure 5) (Fortuna et al., 2017; Maréchal and Plaghki, 1979; Meijer et al., 1998; Power et al., 2014a). rFD is proportional to work of shortening (i.e., the product of force and displacement) such that greater forces or shortening amplitudes increase the magnitude of rFD (Chen et al., 2020; Hahn et al., 2023; Herzog et al., 2000) likely owing to actin angular deformation inhibiting cross-bridge attachment (Herzog et al., 2000; Josephson and Stokes, 1999; Joumaa et al., 2012; Joumaa et al., 2015). Instantaneous stiffness was decreased in both young and old following active shortening as compared with the fixed-end isometric reference contractions, and this is consistent with previous findings that stiffness decreases in proportion to the magnitude of rFD, indicating a decreased proportion of attached cross-bridges (Lee and Herzog, 2003; Power et al., 2014a; Sugi and Tsuchiya, 1988). Our rFD values in both young and old were tightly coupled with stiffness depression (%) (Figure 6, R^2^=0.6-0.8) which aligns with findings in young humans, rats, and rabbits (Joumaa et al., 2012; Joumaa et al., 2015; Mashouri et al., 2021; Pinnell et al., 2019).

Single muscle fibres from the old group performed less mechanical work during active shortening (Figure 5), and this is likely owing to age-related impairments in force production, but also a general slowing of maximal shortening velocity in old age (Hinks et al., 2024; Power et al., 2013b; Thom et al., 2007). Therefore, the absolute shortening speeds used in the present study likely represented a relatively higher percentage of maximal shortening velocity in old as compared with young. Based on the force-velocity relationship, this further disadvantaged fibres from old rats for force production during a shortening contraction ultimately resulting in impaired ability to generate work (Joumaa et al., 2015). Interestingly, despite performing less mechanical work during the shortening contractions than young, the old group experienced considerably greater rFD and stiffness depression (Figure 5). Therefore, age-related changes to the structure and function of the contractile machinery likely altered this history-dependent property of muscle.

### Why did old incur a greater magnitude of rFD than young?

Based on the greater stiffness depression in old as compared with young (Figure 5), there are fewer attached force producing cross-bridges during the isometric steady-state following active shortening in old as compared with young. It is especially likely that greater stiffness depression contributed to the greater rFD in old age considering there were identical magnitudes of increase (38%) in both stiffness depression and rFD in old compared to young, with marked relationships between rFD and stiffness depression. Old, however, had weaker relationships between stiffness depression and rFD (R^2^=0.6-0.65) compared to young (R^2^=0.78-0.84), so other factors besides greater stiffness depression may have contributed to the age-related differences in single fibre rFD. For example, of the remaining attached cross-bridges in the rFD isometric steady-state, there may be more cross-bridges in a weakly bound configuration in old as compared with young (Frontera and Ochala, 2015; Ochala et al., 2007), which could enhance the magnitude of rFD. Further investigations of structural changes during and following active shortening are warranted to determine if shortening-induced actin deformation is greater in old than young.

## Conclusion

We showed that rFD was amplified at the cellular level in old rat single muscle fibres owing to a greater inhibition of cross-bridge attachment as compared with young, while rFE is unaltered. While specific force (i.e., muscle quality) was not different between young and old, there are underlying age-related modifications to the contractile machinery which led to alterations to rFD in old age.

## Acknowledgements

This research was made possible in part using rodents from the National Institute on Aging (NIA). We thank Emma F. Hubbard, Ben Dalton, and Parastoo Mashouri for assistance with some of the initial data collection, and K. Josh Briar for assistance with some of the data analysis.

## Ethics statement

All procedures were approved by the Animal care committee of the University of Guelph (AUP: #4905).

## Data accessibility

Supporting data are available upon request.

## Authors’ contributions

BSN, AH, and GAP conceived and designed the research. BSN, MAP, BED, EFH performed experiments. BSN, AH and GAP analyzed data, interpreted results, prepared figures, and drafted the manuscript. All authors edited and revised the manuscript. All authors have read and approved the final version of the manuscript and agree with the data and order of presentation of the authors.

## References

Abbott, B. C. and Aubert, X. M. (1952). The force exerted by active striated muscle during and after change of length. J. Physiol. 117, 77–86.

Bakenecker, P., Weingarten, T., Hahn, D. and Raiteri, B. (2022). Residual force enhancement is affected more by quadriceps muscle length than stretch amplitude. eLife 11, e77553.

Bottinelli, R., Canepari, M., Pellegrino, M. A. and Reggiani, C. (1996). Force-velocity properties of human skeletal muscle fibres: myosin heavy chain isoform and temperature dependence. J. Physiol. 495 (Pt 2), 573–586.

Burkholder, T. J. and Lieber, R. L. (2001). Sarcomere length range during animal locomotion. J. Exp. Biol. 204, 1529–1536.

Chapman, N., Whitting, J., Broadbent, S., Crowley-McHattan, Z. and Meir, R. (2018). Residual Force Enhancement in Humans: A Systematic Review. J. Appl. Biomech. 34, 240–248.

Chen, J., Hahn, D. and Power, G. A. (2019). Shortening-induced residual force depression in humans. J. Appl. Physiol. 126, 1066–1073.

Chen, J., Mashouri, P., Fontyn, S., Valvano, M., Elliott-Mohamed, S., Noonan, A. M., Brown, S. H. M. and Power, G. A. (2020). The influence of training-induced sarcomerogenesis on the history dependence of force. J. Exp. Biol. 223, jeb218776.

Edman, K. A., Elzinga, G. and Noble, M. I. (1978). Enhancement of mechanical performance by stretch during tetanic contractions of vertebrate skeletal muscle fibres. J. Physiol. 281, 139–155.

Fortuna, R., Groeber, M., Seiberl, W., Power, G. A. and Herzog, W. (2017). Shortening-induced force depression is modulated in a time- and speed-dependent manner following a stretch-shortening cycle. Physiol. Rep. 5, e13279.

Frontera, W. R. and Ochala, J. (2015). Skeletal Muscle: A Brief Review of Structure and Function. Calcif. Tissue Int. 96, 183–195.

Fukutani, A., Misaki, J. and Isaka, T. (2017). Influence of Joint Angle on Residual Force Enhancement in Human Plantar Flexors. Front. Physiol. 8, 234.

Hahn, D., Han, S. and Joumaa, V. (2023). The history-dependent features of muscle force production: A challenge to the cross-bridge theory and their functional implications. J. Biomech. 152, 111579.

Hämäläinen, N. and Pette, D. (1993). The histochemical profiles of fast fiber types IIB, IID, and IIA in skeletal muscles of mouse, rat, and rabbit. J. Histochem. Cytochem. Off. J. Histochem. Soc. 41, 733–743.

Hepple, R. T. and Rice, C. L. (2016). Innervation and neuromuscular control in ageing skeletal muscle. J. Physiol. 594, 1965–1978.

Herzog, W. (2004). History dependence of skeletal muscle force production: Implications for movement control. Hum. Mov. Sci. 23, 591–604.

Herzog, W. (2018). The multiple roles of titin in muscle contraction and force production. Biophys. Rev. 10, 1187–1199.

Herzog, W. and Leonard, T. R. (2002). Force enhancement following stretching of skeletal muscle: a new mechanism. J. Exp. Biol. 205, 1275–1283.

Herzog, W., Leonard, T. R. and Wu, J. Z. (2000). The relationship between force depression following shortening and mechanical work in skeletal muscle. J. Biomech. 33, 659–668.

Hessel, A. L., Kuehn, M., Palmer, B. M., Nissen, D., Mishra, D., Joumaa, V., Freundt, J., Ma, W., Nishikawa, K. C., Irving, T., et al. (2023). The distinctive mechanical and structural signatures of residual force enhancement in myofibers. BioRxiv Prepr. Serv. Biol. 2023.02.19.529125.

Hinks, A., Patterson, M. A., Njai, B. S. and Power, G. A. (2024). Age-related blunting of serial sarcomerogenesis and mechanical adaptations following 4 weeks of maximal eccentric resistance training. J. Appl. Physiol. Bethesda Md 1985.

Hubbard, E. F., Mashouri, P., Pyle, W. G. and Power, G. A. (2023). The effect of gradual ovarian failure on dynamic muscle function and the role of high-intensity interval training on mitigating impairments. Am. J. Physiol. Cell Physiol. 325, C1031–C1045.

Josephson, R. K. and Stokes, D. R. (1999). Work-dependent deactivation of a crustacean muscle. J. Exp. Biol. 202, 2551–2565.

Joumaa, V., Leonard, T. R. and Herzog, W. (2008). Residual force enhancement in myofibrils and sarcomeres. Proc. R. Soc. B Biol. Sci. 275, 1411–1419.

Joumaa, V., Macintosh, B. R. and Herzog, W. (2012). New insights into force depression in skeletal muscle. J. Exp. Biol. 215, 2135–2140.

Joumaa, V., Power, G. A., Hisey, B., Caicedo, A., Stutz, J. and Herzog, W. (2015). Effects of fiber type on force depression after active shortening in skeletal muscle. J. Biomech. 48, 1687–1692.

Joumaa, V., Fitzowich, A. and Herzog, W. (2017). Energy cost of isometric force production after active shortening in skinned muscle fibres. J. Exp. Biol. 220, 1509–1515.

Joumaa, V., Smith, I. C., Fukutani, A., Leonard, T. R., Ma, W., Mijailovich, S. M., Irving, T. C. and Herzog, W. (2021). Effect of Active Lengthening and Shortening on Small-Angle X-ray Reflections in Skinned Skeletal Muscle Fibres. Int. J. Mol. Sci. 22, 8526.

Kanda, K. and Hashizume, K. (1989). Changes in properties of the medial gastrocnemius motor units in aging rats. J. Neurophysiol. 61, 737–746.

Ledvina, M. A. and Segal, S. S. (1995). Sarcomere length and capillary curvature of rat hindlimb muscles in vivo. J. Appl. Physiol. Bethesda Md 1985 78, 2047–2051.

Lee, H.-D. and Herzog, W. (2003). Force depression following muscle shortening of voluntarily activated and electrically stimulated human adductor pollicis. J. Physiol. 551, 993–1003.

Leonard, T. R., DuVall, M. and Herzog, W. (2010). Force enhancement following stretch in a single sarcomere. Am. J. Physiol.-Cell Physiol. 299, C1398–C1401.

Lim, J.-Y. and Frontera, W. R. (2022). Single skeletal muscle fiber mechanical properties: a muscle quality biomarker of human aging. Eur. J. Appl. Physiol. 122, 1383–1395.

Lim, J.-Y. and Frontera, W. R. (2023). Skeletal muscle aging and sarcopenia: Perspectives from mechanical studies of single permeabilized muscle fibers. J. Biomech. 152, 111559.

Maréchal, G. and Plaghki, L. (1979). The deficit of the isometric tetanic tension redeveloped after a release of frog muscle at a constant velocity. J. Gen. Physiol. 73, 453–467.

Mashouri, P., Chen, J., Noonan, A. M., Brown, S. H. M. and Power, G. A. (2021). Modifiability of residual force depression in single muscle fibers following uphill and downhill training in rats. Physiol. Rep. 9,.

Mazara, N., Zwambag, D. P., Noonan, A. M., Weersink, E., Brown, S. H. M. and Power, G. A. (2021). Rate of force development is Ca2+-dependent and influenced by Ca2+- sensitivity in human single muscle fibres from older adults. Exp. Gerontol. 150, 111348.

Meijer, K., Grootenboer, H. J., Koopman, H. F. J. M. and Huijing, P. a. J. B. M. (1998). Shortening history effect in maximally and submaximally stimulated muscle. In Proceedings third world congress biomechanics, p. 90.

Narici, M. V., Maganaris, C. N., Reeves, N. D. and Capodaglio, P. (2003). Effect of aging on human muscle architecture. J. Appl. Physiol. 95, 2229–2234.

Ochala, J., Frontera, W. R., Dorer, D. J., Van Hoecke, J. and Krivickas, L. S. (2007). Single skeletal muscle fiber elastic and contractile characteristics in young and older men. J. Gerontol. A. Biol. Sci. Med. Sci. 62, 375–381.

Patterson, M. A., Hinks, A., Njai, B. S., Dalton, B. E., Hubbard, E. F. and Power, G. A. (2024). Stretch-shortening cycles protect against the age-related loss of power generation in rat single muscle fibres. Exp. Gerontol. 190, 112423.

Pinnell, R. A. M., Mashouri, P., Mazara, N., Weersink, E., Brown, S. H. M. and Power, G. A. (2019). Residual force enhancement and force depression in human single muscle fibres. J. Biomech. 91, 164–169.

Power, G. A., Rice, C. L. and Vandervoort, A. A. (2012). Increased Residual Force Enhancement in Older Adults Is Associated with a Maintenance of Eccentric Strength. PLoS ONE 7, e48044.

Power, G. A., Makrakos, D. P., Rice, C. L. and Vandervoort, A. A. (2013a). Enhanced force production in old age is not a far stretch: an investigation of residual force enhancement and muscle architecture. Physiol. Rep. 1, e00004.

Power, G. A., Dalton, B. H. and Rice, C. L. (2013b). Human neuromuscular structure and function in old age: A brief review. J. Sport Health Sci. 2, 215–226.

Power, G. A., Makrakos, D. P., Stevens, D. E., Herzog, W., Rice, C. L. and Vandervoort, A. A. (2014a). Shortening-induced torque depression in old men: Implications for age-related power loss. Exp. Gerontol. 57, 75–80.

Power, G. A., Herzog, W. and Rice, C. L. (2014b). Decay of force transients following active stretch is slower in older than young men: Support for a structural mechanism contributing to residual force enhancement in old age. J. Biomech. 47, 3423–3427.

Power, G. A., Makrakos, D. P., Stevens, D. E., Rice, C. L. and Vandervoort, A. A. (2015). Velocity dependence of eccentric strength in young and old men: the need for speed! *Appl*. Physiol. Nutr. Metab. Physiol. Appl. Nutr. Metab. 40, 703–710.

Power, G. A., Crooks, S., Fletcher, J. R., Macintosh, B. R. and Herzog, W. (2021). Age-related reductions in the number of serial sarcomeres contribute to shorter fascicle lengths but not elevated passive tension. J Exp Biol 224, jeb242172.

Ramsey, K. A., Bakker, A. J. and Pinniger, G. J. (2010). Fiber-type dependence of stretch-induced force enhancement in rat skeletal muscle. Muscle Nerve 42, 769–777.

Ranatunga, K. W. (1982). Temperature-dependence of shortening velocity and rate of isometric tension development in rat skeletal muscle. J. Physiol. 329, 465–483.

Ranatunga, K. W. (1984). The force-velocity relation of rat fast-and slow-twitch muscles examined at different temperatures. J. Physiol. 351, 517–529.

Rassier, D. E. and Herzog, W. (2004). Considerations on the history dependence of muscle contraction. J. Appl. Physiol. 96, 419–427.

Schilling, N., Arnold, D., Wagner, H. and Fischer, M. S. (2005). Evolutionary aspects and muscular properties of the trunk--implications for human low back pain. Pathophysiol. Off. J. Int. Soc. Pathophysiol. 12, 233–242.

Seiberl, W., Power, G. and Hahn, D. (2015). Residual force enhancement in humans: Current evidence and unresolved issues. J. Electromyogr. Kinesiol. 25, 571–580.

Sugi, H. and Tsuchiya, T. (1988). Stiffness changes during enhancement and deficit of isometric force by slow length changes in frog skeletal muscle fibres. J. Physiol. 407, 215–229.

Thom, J. M., Morse, C. I., Birch, K. M. and Narici, M. V. (2007). Influence of muscle architecture on the torque and power–velocity characteristics of young and elderly men. Eur. J. Appl. Physiol. 100, 613–619.

Tomalka, A., Rode, C., Schumacher, J. and Siebert, T. (2017). The active force-length relationship is invisible during extensive eccentric contractions in skinned skeletal muscle fibres. Proc. Biol. Sci. 284, 20162497.

Tomalka, A., Weidner, S., Hahn, D., Seiberl, W. and Siebert, T. (2021). Power Amplification Increases With Contraction Velocity During Stretch-Shortening Cycles of Skinned Muscle Fibers. Front. Physiol. 12, 644981.

Vlahovic, H., Bazdaric, K., Marijancic, V., Soic-Vranic, T., Malnar, D. and Arbanas, J. (2017). Segmental fibre type composition of the rat iliopsoas muscle. J. Anat. 230, 542–548.

